# Drop of Prevalence after Population Expansion: A lower prevalence for recessive disorders in a random mating population is a transient phenomenon during and after a growth phase

**DOI:** 10.1101/2021.09.29.462290

**Authors:** Luis La Rocca, Julia Frank, Heidi Beate Bentzen, Jean-Tori Pantel, Konrad Gerischer, Anton Bovier, Peter M. Krawitz

**Author notes:** These authors contributed equally to this work (Luis La Rocca, Julia Frank).

## Abstract

Despite increasing data from population-wide sequencing studies, the risk for recessive disorders in consanguineous partnerships is still heavily debated. An important aspect that has not sufficiently been investigated theoretically, is the influence of inbreeding on mutation load and incidence rates when the population sizes change. We therefore developed a model to study these dynamics for a wide range of growth and mating conditions. In the phase of population expansion and shortly afterwards, our simulations show that there is a drop of diseased individuals at the expense of an increasing mutation load for random mating, while both parameters remain almost constant in highly consanguineous partnerships. This explains the empirical observation in present times that a high degree of consanguinity is associated with an increased risk of autosomal recessive disorders. However, it also states that the higher frequency of severe recessive disorders with developmental delay in inbred populations is a transient phenomenon before a mutation-selection balance is reached again.

**Author summary:** What determines the recessive disease burden? The empirical observation that the proportion of intellectual disability of autosomal recessive cause is usually higher in offspring of consanguineous partnerships may lead to a misunderstanding about the mechanisms at work. In any population, selection removes pathogenic alleles from the gene pool while the *de novo* mutation rate adds novel pathogenic alleles. For comparable mutation rates, the incidence of severe recessive disease should be comparable in mutation-selection balance, regardless of the mating scheme. Different incidences can therefore only be explained by population dynamics that are not in equilibrium. We studied in simulated populations the time scales in which mutation-selection balance is reached after a growth phase and found that the mating scheme has a big impact on this lag time. When cousins mate preferentially with cousins, a few generations after a ten-fold increase in size the new equilibrium is established. In contrast, for random-mating, the transient advantage of a lower incidence may last for hundreds of generations, while the mutation load increases. By this means, our findings also highlight the importance of better carrier screens in the future for genetic consultations.

## Introduction

The analysis of sequencing data of large cohorts allowed to confirm estimates for the number of recessive, lethal equivalents per genome, that were previously based on epidemiological data of disease prevalences and stillbirths: On average, healthy individuals carry 0.5 to 2 heterozygous variants that would prevent reproduction if they occurred in a homozygous state [1–4]. However, the question of how the ethnic background and the degree of consanguinity affect the recessive lethal load per person is still vividly discussed since empirical data and predictions of theoretical population genetics are partially contradictory [5]: assuming mutation-selection balance the prevalence of recessive disorders should be the same regardless of ethnicity and mating scheme. However, in the DDD cohort, the proportion of cases due to recessive coding variants was 3.6% in patients of European ancestry, compared to 31% in patients with Pakistani ancestry [6]. Even within the same population, e.g. in Iran, the probability for a recessive cause of intellectual disability is four times higher for offspring from first-cousin unions than for offspring of non-consanguineous partnerships [7–9]. To explain this discrepancy between mutation load and recessive disease burden, some authors recently argued that the unexpectedly high frequency of lethal equivalents might also be explained by an ascertainment bias, that is, some of the pathogenic mutations reached high frequency by chance and are therefore overreported [10]. Since also the assumption of mutation-selection balance is flawed, other authors studied the effect of different demographic dynamics including explosive population growth on mutation load [11]. In our work, we explore the influence of different mating schemes on these dynamics by means of two different simulation frameworks. Each model had the advantage of handling certain aspects of population genetics particularly well. The first is an adaption of the classical Wright-Fischer model with discrete non-overlapping generations run in the forward genetic simulation framework SLiM [12, 13]. In the second model, generations can overlap, because diploid individuals die and give birth at independent exponential times on a continuous timescale [10]. Interestingly, simulations of the discrete, as well as the continuous model yield the same results, which suggest that the lower recessive disease burden in outbred populations is only a transient advantage at the expense of an increasing mutation load in comparison to an inbred population.

## Results

In this paper we investigate the dynamics of the average number of lethal equivalents per genome (mutation load) and the recessive disease burden in the population (prevalence) during and after a growth phase for different mating schemes. We assess mutation load and prevalence in a discrete and a continuous simulation framework in dependence of the generations, respectively the time that passed. In the following, we describe the behaviour of these variables for a population that starts with 500 individuals and grows logistically to a size of 10 000. Later on, we also discuss the effect of different genomic architectures, effective population sizes and mutation rates, which yield similar a dynamic behavior for prevalence and mutation load.

At the beginning of the simulation the population consists of genetically identical individuals. Their diploid genomes are composed of 1 000 recessive genes that are crucial for reproductive success. The length of the coding sequences per gene ranges between 500 and 10 000 bp and novel alleles are introduced with a *de novo* mutation rate of 1.2 · 10^−8^ per bp per offspring. One out of nine such mutations is pathogenic, while the others are considered neutral. The choice of these parameters is motivated by the distribution of coding lengths and the deleteriousness scores of known autosomal recessive genes [14, 15]. Whenever both copies of a gene carry at least one pathogenic allele, the individual is *affected* and cannot propagate. All pathogenic mutations are therefore lethal equivalents in our simulations.

The two mating schemes that we study differ in the choice of partners. When mating partners are chosen randomly, this results in an outbred population, while mating within the same family results in an inbred population. We implemented the later scheme by enforcing 70% of partnerships between cousins or individuals of comparable relatedness which is representative for populations with a high prevalence of consanguinity (consang.net).

During an initial burn-in phase of 500 generations a mutation-selection balance is reached for both mating schemes. That means, the mean number of pathogenic alleles that enter the gene pool by *de novo* events equals the mean number of pathogenic alleles that are removed due to affected individuals that do not propagate. The neutral alleles do not have any effect on fitness and rather serve as a control for known findings from the literature. For instance, in equilibrium the allele frequency distribution of such variants is predicted by Kimura’s theory of neutral molecular evolution.

At generation 500 the population starts to grow. In the discrete model this is simulated by a deterministic logistic function, and the birth and death rates for the continuous time model are set to mimic this growth by a rate of *r* = 10^−5^. After roughly 130 generations the growth phase ends as the population has reached its new target size of 10 000 individuals. As we will see in the following, both prevalence and mutation load require many more generations to reach their new equilibria. Therefore, although the growth curve is logistic, it rather looks like a step function on a scale of 2 000 generations (grey curve in Fig. 1). The most remarkable change in the random mating population is the almost instantaneous drop in prevalence when the expansion starts. Shortly after the end of the growth phase, the prevalence has reached its lowest point, which is down by two thirds. From there it starts increasing again, but more slowly, until it reaches the initial level of 0.7% approximately 750 generations after the start of the expansion. Likewise, the mutation load increases very fast when the population starts to grow. Roughly 550 generations after the population has reached its new equilibrium the mutation load has quadruplet and approximates a new plateau at three deleterious mutations in 1 000 recessive disease genes.

**Fig. 1.**
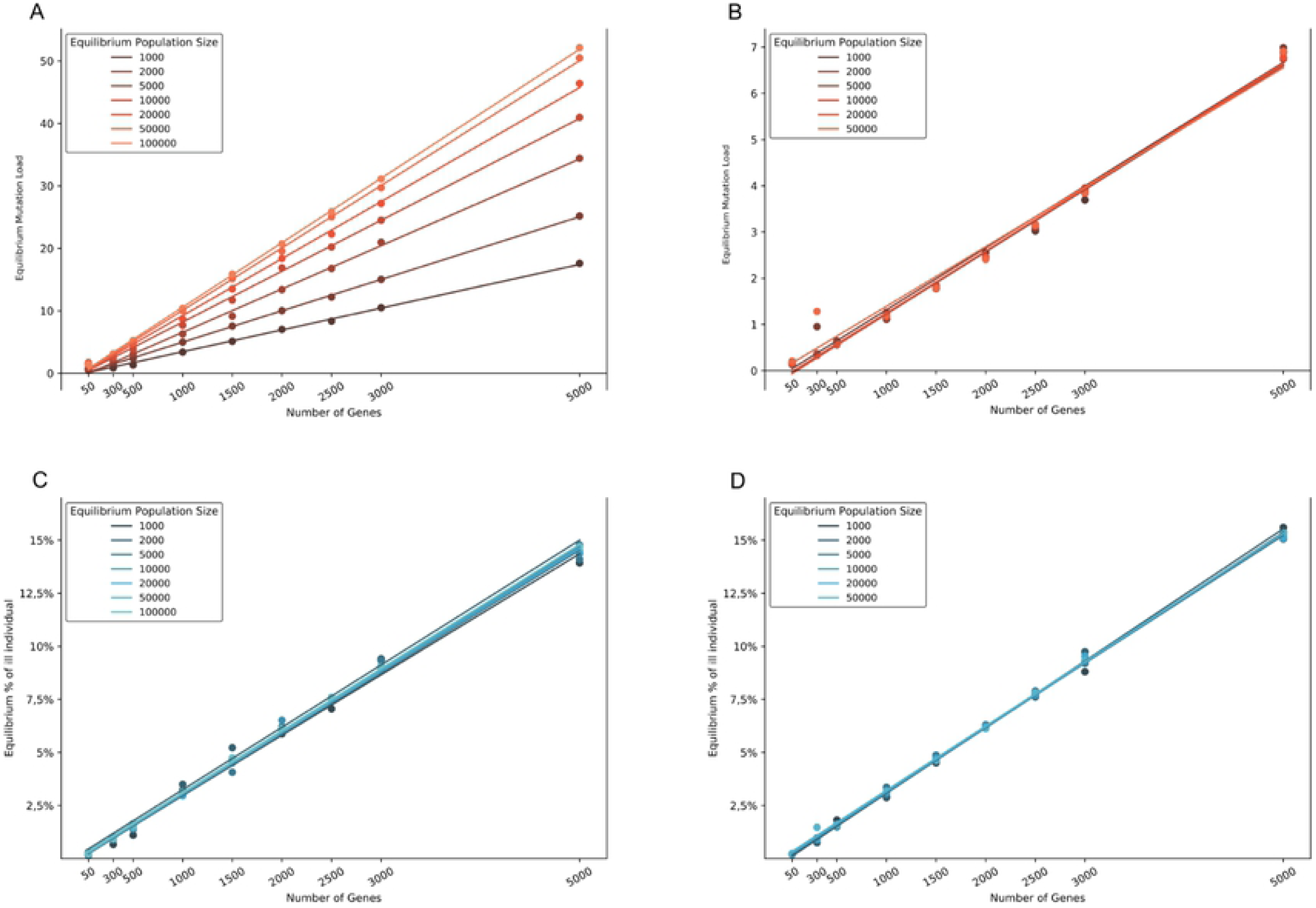
Dynamics of mutation load and prevalence for severe recessive disorders. A population expansion from 500 to 10 000 individuals (grey), starting in generation 500 does not affect prevalence (orange) nor mutation load (red) if partners are preferentially chosen within relatives (consanguineous mating scheme) (A). In contrast, in a random mating population, there is a transient drop of prevalence at the expense of an increasing mutation load (B). It takes more than 550 generations after the end of the growth phase, until the steady state is reached and the prevalence for both mating schemes are comparable again. The plots show the average of 50 exact trajectories of the stochastic process simulated with the discrete model.

In contrast, the prevalence and mutation load in the inbred population remain almost constant, and only the variance is reduced due to the larger population size. The prevalence stays at the level of roughly 70 affected individuals per generation in a population of 10 000 whereas the mutation load maintains its low level of 0.25 mutations per individual during the whole time of nearly 2000 generations. This effect is best explained by the effective number of potential partners for an individual within both mating schemes. In a randomly mating population the number of potential partners increases considerably with the size of the population, whereas in a consanguineous population this number stays almost constant during the expansion, because most individuals are forced to mate within the same family. On the other hand, the individuals that mate outside the family give rise to new clans, explaining the population growth.

We also studied the equilibrium mutation load and prevalence as a function of mutation rate and genome architecture ((Fig. 3)). In mutation-selection balance, prevalence and mutation load depend only on the mutation rate and the genomic architecture but not the mating scheme. For a single gene architecture, the square root of the mutation rate determines the frequency of lethal equivalents. Higher mutation rates will result in higher prevalence and mutation load. Likewise, increasing the number of recessive genes, while keeping the mutation rate constant will increase mutation load and prevalence. The mutation load will also increase with population size in a random mating scenario, in fact the faster the more recessive genes there are. In contrast, in an highly inbred population, its effective size is bound. Therefore, the equilibrium mutation load will also depend on the average size of a family or clan.

Finally, we simulated the dynamics of mutation load and prevalence in a consanguineous population consisting of clan sizes from 10 individuals up to 10,000 individuals (Fig. 2). The larger the clans become, the more the dynamics resemble random mating. Already with a clan size of 100 individuals a small drop of diseased individuals is observable when the population starts growing. For larger clan sizes also the equilibrium mutation load becomes significantly higher.

**Fig. 2.**
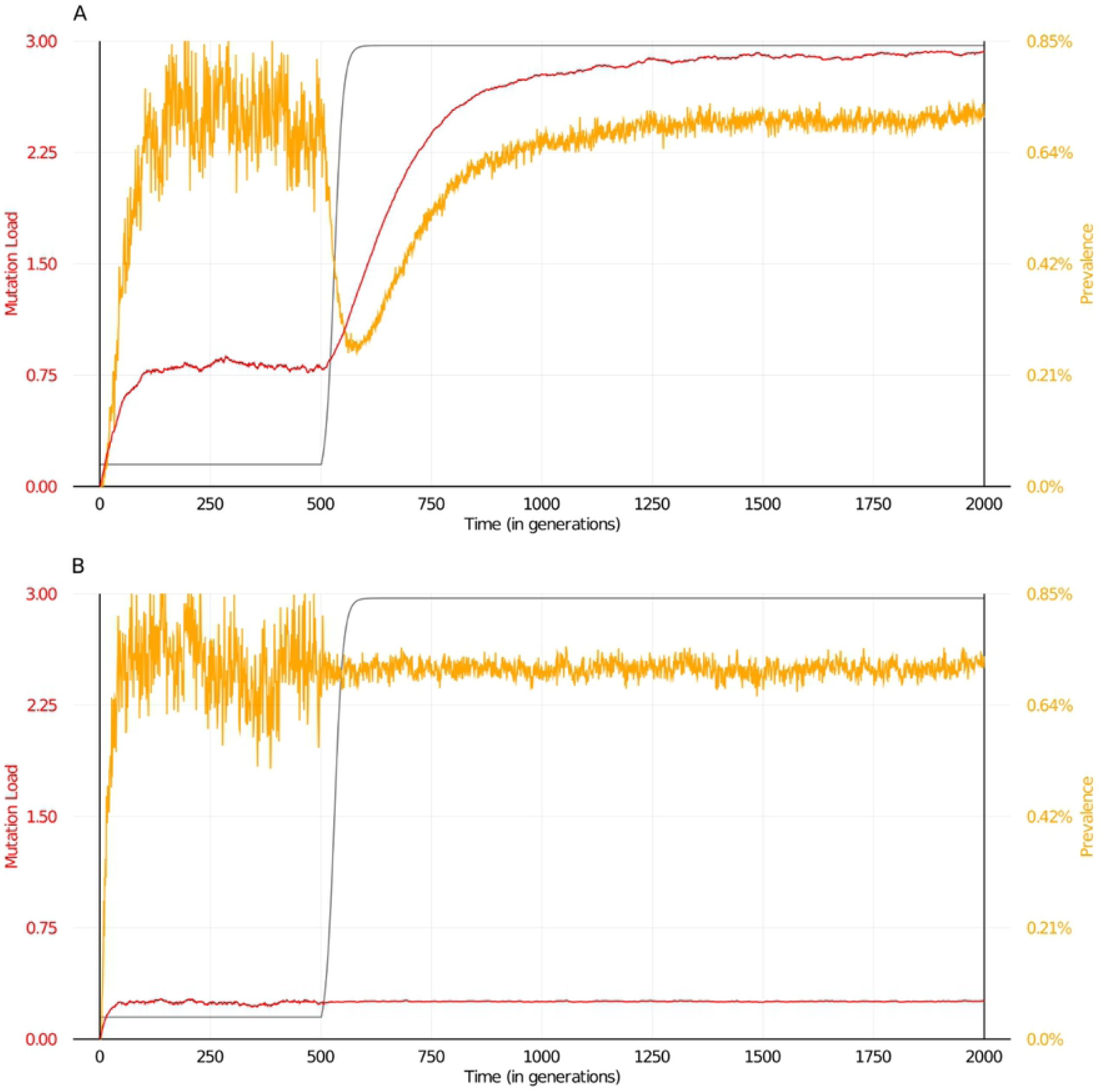
Influence of family size on mutation load and prevalence. The mating scheme is characterized by the family size and a probability function that describes how many of the partners are chosen within the family. In a preferentially consanguineous mating population the dynamics change when the maximum family size increases (upper left panel to lower right from 10, 25, 50, 100, 500 up to 10 000). The mutation load starts to increase considerably if mating is happening in tribes of 500 individuals. However, at this stage there is still only a minor effect of further population growth. In the lower right the maximum of the allowed family size is equivalent to the population size and thus, dynamics do not differ from a random mating scheme any more. The plots show the average of 10 exact trajectories of the stochastic process simulated with the continuous model.

**Fig. 3.**
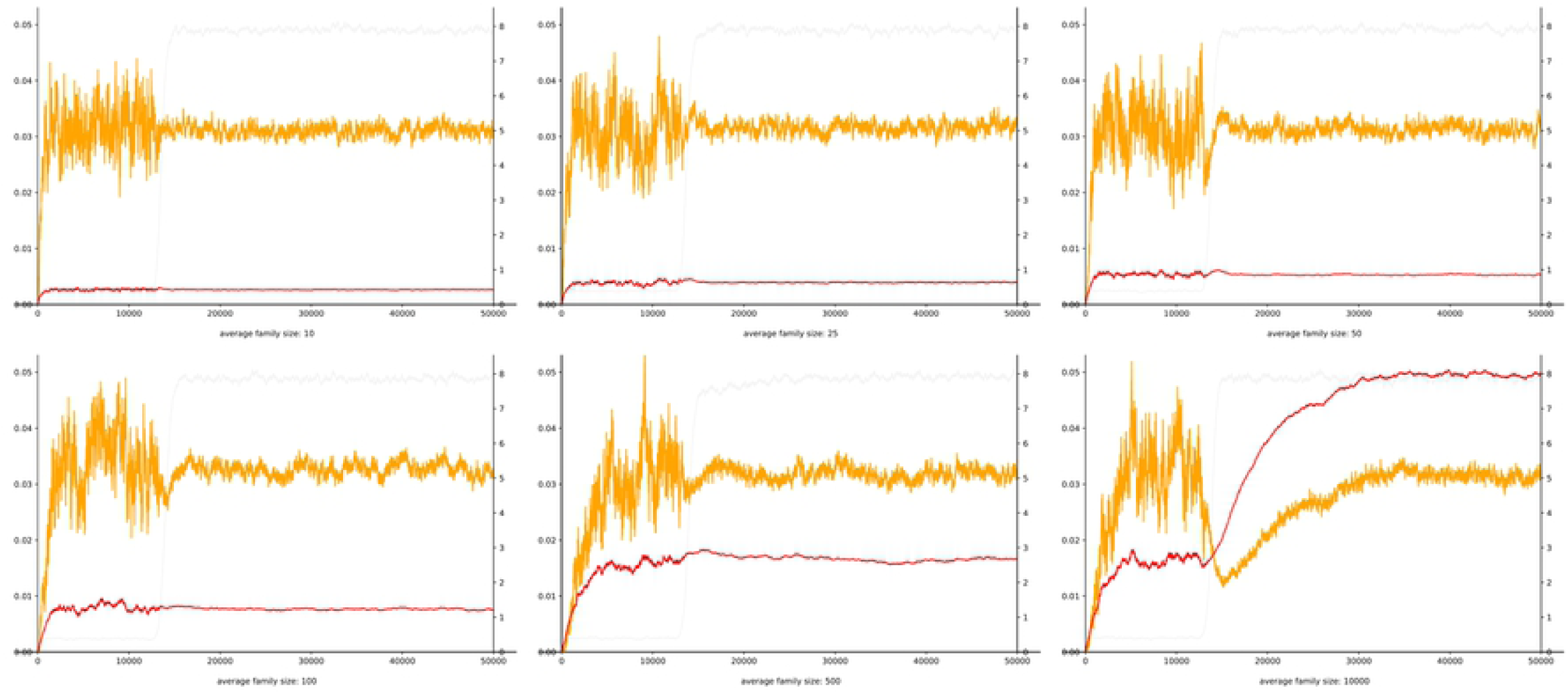
Influence of genomic architecture and population size. The capacity of the genome for deleterious mutations is larger in the random mating population. With an increasing number of genes and growing population size, deleterious mutations accumulate (A). In contrast, in the consanguineous mating scheme, family size limits the effective population size, and therefore mutation load is independent of the total number of individuals (B). Prevalence increases linearly in both mating schemes when the number of genes increases and is independent from population size, as regression analysis indicates (C,D).

Since the change in mutation load corresponds to the transient reduction of prevalence, the dynamics are reversed, when a large population has to go through a bottleneck. Reducing the population is equivalent to higher inbreeding and the mutation load will decrease after a spike in prevalence. Thus, whether the transient effects on prevalence in a random mating population are beneficial or detrimental depends critically on demographics.

## Materials and methods

In both simulation frameworks, the mutation load is defined as the average number of lethal equivalents per individual. The lethal equivalents in the genome are deleterious alleles that are disease-causing if both copies of a gene in an individual harbor at least one such variant. The totality of these mutations could also be regarded as the theoretical super-set for an extended carrier screen [16]. Note that the this definition of mutation load does not match the classical definition, where differences in fitness are decisive [17]. Here all individuals are equally fit independent of their mutation load unless they do not carry a lethal equivalent on both copies of a gene. In that case their fitness decreases to zero. By this means, we are able to focus on the incidence rate of severe recessive disorders with early onset that prevent reproduction almost with certainty. Likewise, we can study how the selection of a partner, which we refer to as a mating scheme, influences the disease prevalence and mutation load and we are able to monitor these parameters in the population over time. This is done by counting the number of lethal equivalents that enter the gene pool due to a constant *de novo* mutation rate, or leave the gene pool due to selection. If the disease prevalence does not change any more, the population is in a steady state, that is a flux balance for lethal equivalents.

In our simulations, affected individuals do not have a different life span from unaffected individuals, therefore prevalence and incidence are equivalent and their rate is proportional to the amount of lethal equivalents removed from the gene pool per generation. In fact, the expected number of lethal equivalents that is lost by an affected individual that is not propagating is two. This is equivalent to the difference in the average mutation load between affected and unaffected individuals and can also be derived from the simulations. An expansion of the population will affect prevalence and mutation load as we will discuss in more detail in the following.

Consider a finite population of individuals where each individual is characterized by a diploid set of *N* gene segments of different size. Mutations appear at every gene independently with a rate that is proportional to its size. As long as an individual carries a deleterious mutation at only one gene its fitness is unaffected. But as soon as both copies of a gene carry a mutation the individuals reproductive fitness is reduced to zero. Hence it will be excluded from the mating process and is not able to reproduce anymore. One can think of a mutation causing a severe recessive disorder. Besides that all individuals are equally fit, no matter how many recessive disorders they carry.

Simulations always start with a small, healthy population. After a period of time in which a mutation selection balance is established a logistic growth phase starts, that settles after a new target size for the population is reached. We investigate changes of the dynamics of the mutation load and the prevalence when the population applies different mating schemes. On the one hand random mating, with an equal probability for each two individuals with non-zero fitness to mate. And on the other hand a consanguineous mating scheme, where individuals prefer to mate with close relatives.

### Discrete Model

In the default setting, the simulation package from Haller and Messer [12] samples a diploid population evolution according to the standard Wright Fisher model. Sexes were added such that each sex is equally represented in the population at any time. In generation *n* ≥ 1 there is a finite number of individuals *M*_*n*_ ≥ 0 with a total of 2*M*_*n*_ genomes alive. In the initial burn-in phase the population size is held constant such that *M*_*n*_ = *M*_0_ for all generations *n* ≤ *n*_*grow*_. Afterwards the growth phase begins and the population size of each generation grows logistically with growth rate *r* > 0 until it approaches the new target size *K*. Therefore the population size of each generation is determined by the following formula

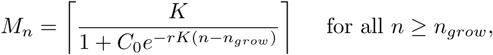

where 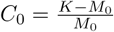

The two mating schemes - random and consanguineous mating - are introduced as following. To create generation (*n* + 1) first choose *M*_*n*+1_ women from generation *n* independently at random with replacement among all women with non-zero fitness. For random mating each woman then selects a man uniformly at random from the pool of potential partners with truly positive fitness. To implement the consanguineous mating scheme use the pedigree information SLiM provides for the last two generations backwards in time. Thus for every individual we know who are its parents and its grandparents. A woman in the consanguineous population now chooses a mate with a weighted uniform distribution on the set of all potential partners. For some weights *α, β* ∈ [0, 1] with *α* + *β* ≤ 1 the individual chooses a man with

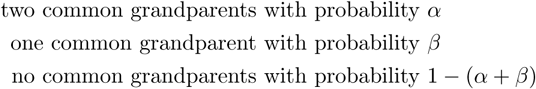

Notice that having two grandparents is similar to being cousins and having one grandparent in common relates to being half-cousins.

From the whole human genome pick *N* gene segments with an independent, uniformly distributed number of base pairs *w*_1_, …, *w*_*N*_ ∼ 𝒰_[*a,b*]_, where *a, b* > 0 is the minimum resp. maximum segment size. Moreover the whole genome is divided into *n*_*c*_ chromosomes. At birth not only mutation alters the offsprings genetic information, but also recombination. For each chromosome start an independent *Poisson* Process with rate *r*_rec_ > 0 marking the recombination breakpoints. Here *r*_rec_ is the overall recombination rate.

### Continuous Model

The adaptive dynamics model is continuous in time, hence time is not measured in *n* ∈ ℕ discrete generations, but on the positive real axis *t* ∈ ℝ_+_. No exact pedigree are available for this continuous model, therefore introduce a new diploid family trait, which indicates the ancestry of an individual. Hence every individual is characterized by two diploid traits. The first refers to the family origin whereas the second gives insight in the genetical information of the individual. Introduce ℱ ⊂ ℕ as the finite set of all possible family traits and **f** = (*f* ^1^, *f* ^2^) ∈ ℱ^2^ being the family trait of an individual in the current population. Moreover the diploid genetic information of an individual is a finite vector

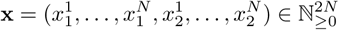

where the entries 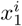 and 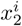 represent the number of mutation at the *i*^th^ gene segment in the paternally and maternally inherited haploid genomes. Thus if there are 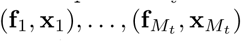 individuals alive at time *t* > 0 in an arbitrary order, define the population state as a point measure on 𝒳 := (ℱ^2^×ℕ^2*N*^)

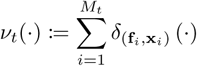

Individuals give birth and die at exponential rates *b*(**f**, **x**) resp. *d*(**f**, **x**) which depend on the family trait and the chromosomal configuration of the individuals. One can think of every individual carrying two independent clocks, a birth and a death clock. If the birth clock rings first the individual makes its mating choice, reproduces and resets its clock. Whereas if the death clock rings the individual disappears from the population.

Additional to the intrinsic death rate every individual senses the competition pressure of every other individual in the population. The term *C*(**f**, **x, g, y**) gives the competition pressure executed by an individual of type (**g, y**) and felt by an individual of type (**f**, **x**). Hence the total death rate of an individual in population *ν* is increased by the term

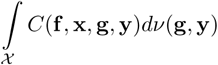

When an individual gives birth it chooses a partner at random from the population according to the partners birth rate and the reproductive compatibility between them. Notice that the reproductive compatibility *R*_**f**_ (**g**) ∈ [0, 1] of two individuals depends only on their family traits **f** and **g**. After a mate was chosen the newborns family trait will be a uniform random combination of the four parental family traits unless both parents have the same traits. In this case the child inherits the exact same couple of traits. The genetic configuration of the newborn is not only a random combination of the parental alleles since the effect of mutation and the reshuffling of the parental chromosomes come into play. For an accurate definition of the reshuffling of the diploid parental chromosomes to a mixed haploid set, define first the sections on the genetic information, that form chromosomes. Let *n*_*c*_ ∈ ℕ be the number of chromosomes for every individual. Introduce the chromosome breakpoints 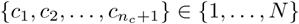 with 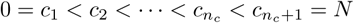. Divide the genetic information of every individual 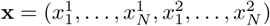 into chromosomes of the same length in both copies

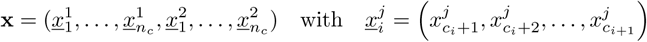

for *j* ∈ {1, 2} and *i* ∈ {1, …, *n*_*c*_}. Finally get the reshuffled gamete from **x** via a selection variable *τ* : {1, 2, …, *n*_*c*_} → {1, 2} as follows

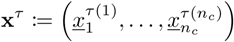

Selecting *τ* among all possible assignments equals an uniform recombination of the diploid chromosomes into haploid set. At each birth a *Poisson* distributed number of mutations is added to every offspring. The expectation of these identically distributed and independent *Poisson* random variables equals the total mutation rate 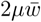, where *µ* is the mutation rate per base pair and 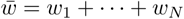 is the sum of the length of the gene segments under consideration. After sampling the number of mutations per birth, these mutations are distributed equally on the 2*N* gene segments according to their length. Finally a mutation at a given gene increases the genetic value by one. All of this is captured in the mutation with recombination operator 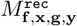 for a mating between individuals (**f**, **x**) and (**g, y**)

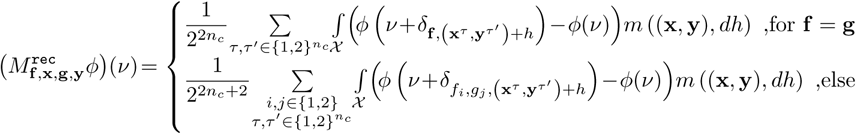

and the mutation measure *m* is defined as

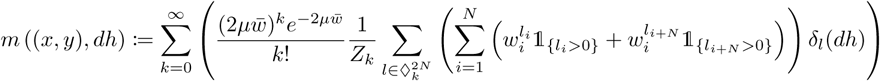

where 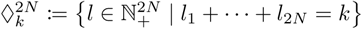 is the set of all lattice vectors in 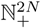 with one norm equal to *k* moreover *Z*_*k*_ > 0 is a normalizing constant depending on the size of 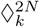. Let

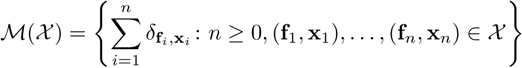

be the set of all finite point measures on 𝒳. The dynamics of the continuous time, ℳ(𝒳)-valued jump process (*ν*_*t*_)_*t*≥0_ can be described by the generator ℒ defined for any bounded measurable function *ϕ* : ℳ(𝒳) → ℝ as

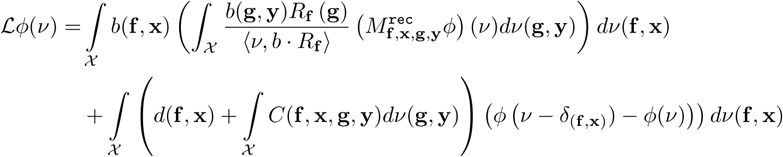

Assume the boundedness of the birth and death rates *b* and *d* as well as the boundedness of the competition kernel *C*. Starting in a initial state *ν*_0_ ∈ 𝒳 such that

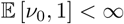

existence and uniqueness in law of the process with infinitesimal generator ℒ and initial condition *ν*_0_ can be derived from [18].

### Keep families small

To ensure a stable mating scheme during the evolution of the population split families which became too big up into two subfamilies. Therefore introduce the following sequence of stopping times.

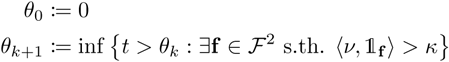

for some fixed *κ* > 0. Notice that at any stopping time it is always a unique family **f** ∈ ℱ^2^ that exceeds the maximum family size, since at any time there is at most one individual entering or exiting the population. At these random times the big family is split up at random into two subfamilies, where the size of each subfamily is binomial distributed with mean 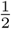. One family keeps the old family trait and the other one gets a completely new, homogeneous one. To make this precise associate a number to each individual in a family. Therefore let 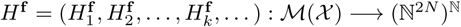 defined by

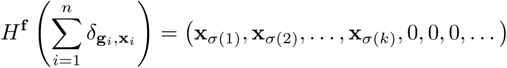

where the **x**_*i*_ for *i* = 1, …, *k* are the genetic configurations of the individuals in ((**g**_1_, **x**_1_), …, (**g**_*n*_, **x**_*n*_)) with **g**_*i*_ = **f** and where **x**_*σ*(1)_ ⪯… ⪯ **x**_*σ*(*k*)_ is the lexicographical order ⪯ on ℕ^2*N*^ and *k* = ⟨*ν*, 𝟙_**f**_ ⟩ is the family size of **f**. Then the splitting of a family with trait **f** ∈ ℱ^2^ can be expressed with the following operator for any bounded measurable function *ϕ* : ℳ_*F*_ (𝒳) →ℝ

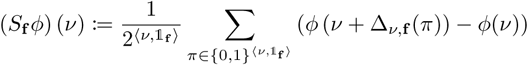

where Δ_*ν*,**f**_ (*π*) executes the splitting of the family **f** in *ν* into two with configuration *π*, hence

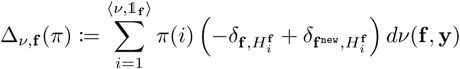

where **f** ^new^ = (*f* ^new^, *f* ^new^) with *f* ^new^ ∈ {*g* ∈ ℱ | ⟨ *ν*, 𝟙_(*g,g*′_) ⟩ = 0, ∀*g*′ ∈ ℱ} chosen deterministically, is a homogeneous family trait which is entirely new to the population. A possible way of choosing the new family trait at time *θ*_*k*_ is to set *f* ^new^ = *n*_ℱ_ + *k* where *n*_ℱ_ is the number of different families in the initial population *ν*_0_. Hence the dynamics of the evolutionary process with splitting is given as the solution of the following martingale problem. Let *ν*_0_ ∈ ℳ(𝒳) be a initial population then for any real, continuous, bounded function *ϕ* on ℳ(𝒳) the process

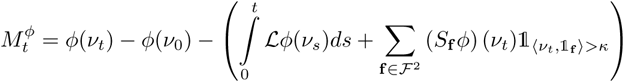

is a martingale.

### Choices of Parameters

Introduce the subset 𝒟_*N*_ ⊆ 𝒳 of traits having at least one mutation in the same gene segment on both copies of the chromosome as

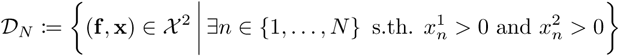

and set the birth and death rate to constant unless an individual falls in the set of non-propagable types

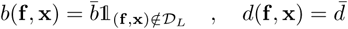

for 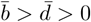. To ensure uniform competition among all individuals set the competition pressure constant to

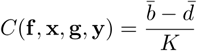

for all individuals (**f**, **x**) and (**g, y**). Therefore the population size in equilibrium fluctuates around the target size *K* of the system. The different mating schemes are defined as follows. First the random mating, where 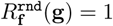 for all **f**, **g** ∈ ℱ^2^ and the consanguineous mating scheme with

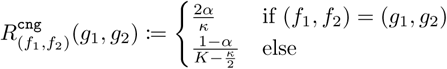

where *α* ∈ [0, 1) and *κ* > 0 is the maximum family size. Therefore the probability of mating within their own family that is of size 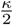 in a population that is at its stable equilibrium size *K* is constant *α*. Note that for families with family size smaller than 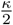 the probability of mating within the family is slightly lower, whereas it gets bigger when the family size surpasses this size. Furthermore the probability increases at the beginning of the growth phase when the carrying capacity *K* gets uplifted and the population size starts to grow slowly. During this initial phase of expansion there will be more consanguineous mating overall. This imbalance levels off as soon as the population size approaches *K*.

Start the evolution with a small, healthy population of size *M*_0_ > 0 in population equilibrium, where nobody carries any mutation. The clonal individuals are following divided into *n*_ℱ_ ∈ ℕ_>0_ families with homogeneous family traits (1, 1), (2, 2), …, (*n*_ℱ_, *n*_ℱ_). After an initial phase during which a mutation selection balance is established raise the carrying capacity to generate a natural exponential population growth up to the new equilibrium. The population parameters we are particular interested in, the mutation load and the prevalence rate for the disability can be formulated in terms of the population process. The relative mutation load of a population *ν* is defined as

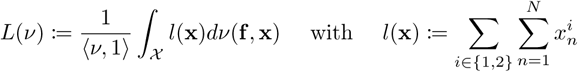

And the relative number of individuals in the population *ν* belonging to the set 𝒟_*N*_ is

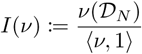

To generate stochastically correct trajectories of the population dynamics process we implemented a variation of Gillespie algorithm for the above model in Python.

### Comparing both models

Both models, the discrete generation model implemented with SLiM and the continuous time model with the Gillespie algorithm, fit different aspects of nature better than the other. A key feature of SLiM and the discrete model is the exact pedigree information generated for every individual. Whereas the continuous model can only cluster roughly in family clans, but cannot differ between members of one family. A major drawback of the discrete model are the non overlapping generations. That excludes the possibility of matings between individuals on different pedigree levels such as uncle - niece marriages. This deficit is set aside by the continuous time model. Since individuals give birth and die independently, all individuals are of different age, and thus different generations are alive simultaneously. On the one hand alike the Wright-Fisher model, the discrete model works with constant respectively deterministically increasing population sizes. On the other hand the continuous model has a fluctuating and natural growing population. Notice that for large populations the random fluctuation in population size are of order 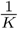 and the stochastic process converges in law to the solution of a deterministic, logistic equation [18].

Recombination is also implemented differently. While SLiM works with true inter chromosomal recombination, the continuous model only reshuffles the parental chromosomes while producing the gamete. This is due to the fact that SLiM saves the exact base position of mutations on the human genome, whereas the continuous model only knows the number of mutations per gene segment and not their exact location within each segment. Since genes are short compared to the entire length of a chromosome and only about two recombinations occur per meiosis, mixing maternal and paternal variants of a gene in the gametes occurs rarely. Furthermore, the effect on fitness is the same for all pathogenic alleles, no matter whether they occur in a homozygous or compound heterozygous state.

Besides all differences parameters are chosen to be equal for both simulations. Among these the number of gene segments *N*, the initial and equilibrium population size *M*_0_ and *K*, and many more. Moreover family sizes in the continuous model are chosen such that the number of potential partners in both models in the consanguineous setting are approximately equal. Likewise the birth rate in the continuous time model is set to *b* = 1, such that in generation *t* there are *M*_*t*+1_ birth events, where *M*_*t*_ is the population size at that time. The only difference is, that for the discrete generation model there are *exactly M*_*t*+1_ birth, whereas in the continuous time model there are *on average* that many births

## Discussion

The empirical observation that consanguinity is often associated with an increased recessive disease burden, has been made in several populations for centuries framing risk perception and influencing many aspects of social norms. However, according to our simulations the advantage of outbreeding is a transient phenomenon for a population that is initially in mutation-selection balance and that starts to grow. The lower prevalence compared to an inbreed population lasts for many generations even after the expansion phase has ended until mutation-selection balance is reached again for a higher mutational load. We found it intriguing that e.g. first-cousin marriage in Europe was banned after several generations of population growth during the Roman empire and considerable migration and admixture [11]. While this continent clearly benefited at that time point from a change of social conventions with respect to the recessive disease burden, the consequences of different mating schemes e.g. on the proportion of congenital malformation are less prominent in populations that were more constant in size over a long period of time [3]. The occurrence and coexistence of different marriage patterns over many centuries can certainly not be understood by population genetics alone since social and economic factors interact with demographics in a complex manner [11]. It is therefore irritating when genetic reasoning of questionable validity is used in the legislature. For instance, the European Court of Human Rights case of Stübing v. Germany concerned consanguine siblings who had four children following consensual intercourse, whereupon both siblings were charged with incest [19]. One of the siblings lodged a complaint, arguing that the legislature violated his right to sexual self-determination, his private and family life. The Court found that 24 out of 44 European States reviewed, criminalized consensual sexual acts between adult siblings, and all prohibited siblings from getting married. The German government argued that the law against incest partly aimed to protect against the significantly increased risk of genetic damage among children from an incestuous relationship. However, regardless of the validity of this argument, motivating a law on avoiding a higher probability of disease can be viewed as eugenic. As the German Ethics Council opined after the judgment, no convincing argument can be derived from there being a risk of genetic damage, concurring with a statement from the German Society of Human Genetics that “The argument that reproduction needs to be thwarted in couples whose children possess an elevated risk for recessively inherited illnesses is an attack on the reproductive freedom of all” [20, 21]. In contrast to the legislature, the predicted recessive disease burden that is communicated in a genetic consultation might influence decisions e.g. about the choice of partners or family planning. Since this risk does not only depend on mating schemes but also on mutation load it is important to measure this parameter as accurately as possible. In our simulations an individual of the outbreed population had in average four times more lethal equivalents than an individual of the inbreed population when mutation-selection balance was reached again many generations after the growth phase ended. Interestingly, these values and the range are comparable to what has also been described in the literature for real populations. With respect to the British subpopulations of Pakistani (PABI) and European (EABI) ancestry in Martin, et al., this could mean that PABI with a considerably higher autozygosity and many first-cousin marriages are closer to mutation-selection balance than EABI. This would imply that the disease prevalence for recessive disorders will remain constant for PABI while it will increase in the following generations for EABI, given the different mating schemes. Based on current ClinVar statistics there are more than 150,000 pathogenic alleles known for recessive genes that cause severe disorders. Data of a recent large German cohort indicates that more than half of the families with offspring with a recessive disease could have benefited from exome analysis of the healthy parents. It is therefore important that the increasing knowledge about pathogenic variants is made available in expanded carrier screens [16].

## Supporting information

All scripts to reproduce our simulation results can be found in the following repository: https://github.com/roccminton/Diploid_Model_Two_Loci.

A video clip of our simulations can be found at: https://youtu.be/5hOgLyRqWPg.

## Acknowledgments

This work was partially supported by the Deutsche Forschungsgemeinschaft (DFG, German Research Foundation) under Germany’s Excellence Strategy GZ 2047/1, Projekt-ID 390685813 and GZ 2151, Project-ID 390873048 and through the Priority Programme 1590 “Probabilistic Structures in Evolution”. We thank Benjamin C. Haller for his very helpful advices on how to get the best out of the forward genetic simulation framework SLiM [12].

